# Egg activation triggers clearance of maternally deposited RNA binding proteins

**DOI:** 10.1101/806968

**Authors:** Michael Zavortink, Lauren N. Rutt, Svetlana Dzitoyeva, Chloe Barrington, Danielle Y. Bilodeau, Miranda Wang, Xiao Xiao Lily Chen, Olivia S. Rissland

## Abstract

The maternal-to-zygotic transition (MZT) is a conserved step in animal development, where control is passed from the maternal genome to the zygotic one. Although the MZT is typically considered from its impact on the transcriptome, we previously found that three maternally deposited *Drosophila* RNA binding proteins (ME31B, Trailer Hitch [TRAL], and Cup) are also cleared during the MZT by unknown mechanisms. Here, we show that these proteins are degraded by the ubiquitin-proteasome system. Kondo, an E2 conjugating enzyme, and the E3 CTLH ligase are required for the destruction of ME31B, TRAL, and Cup. Importantly, despite occurring hours earlier, egg activation establishes the timer for clearance of these proteins by activating the Pan Gu kinase, which in turn stimulates translation of *Kondo* mRNA. In other words, egg activation triggers a series of regulatory events that culminate in the degradation of maternally deposited RNA binding proteins several hours later. Clearance of the maternal protein dowry thus appears to be a coordinated, but as-yet underappreciated, aspect of the MZT.

**HIGHLIGHTS:**

- Degradation of ME31B requires the PNG kinase, but not fertilization
- The ubiquitin-proteasome system degrades ME31B via CTLH E3 ligase and the UBC-E2H/Kondo ubiquitin-conjugating enzyme
- The association of ME31B with the CTLH complex does not require PNG activity
- PNG kinase mediates the translational upregulation of Kondo at egg activation

## INTRODUCTION

Proper embryogenesis is critical for animal development. Many of the earliest events occur prior to the onset of zygotic transcription, and they are instead directed by maternally deposited proteins and messenger RNAs (mRNAs). During the maternal-to-zygotic transition (MZT), genetic control of developmental events changes from these maternally loaded gene products to newly made zygotic ones (Vastenhouw et al., 2019). Thus, the MZT requires both the activation of zygotic transcription (also called zygotic genome activation or ZGA) and clearance of maternal transcripts. Failure to mediate either of these processes is lethal for the embryo (Benoit et al., 2009; Liang et al., 2008).

In contrast to our understanding of the transcriptome during the MZT, much less is known about changes in the proteome. Despite the fact that the maternal dowry of proteins plays key roles in embryogenesis, there are only a handful of examples of cleared maternal proteins (Guven-Ozkan et al., 2008; Hara et al., 2017; Pesin and Orr-Weaver, 2007; Sysoev et al., 2016; Wang et al., 2017; Yang et al., 2016). Recently, we found that three RNA binding proteins (ME31B, Trailer Hitch [TRAL], and Cup) are rapidly degraded during the MZT in *Drosophila melanogaster*, at a time point coinciding with the major wave of zygotic transcription (Wang et al., 2017). ME31B, TRAL, and Cup form a complex that blocks translation initiation (Kinkelin et al., 2012; Nakamura et al., 2004; Nelson et al., 2004; Wilhelm et al., 2003). All three proteins are required for oogenesis, and they appear to bind and repress thousands of deposited maternal mRNAs (Keyes and Spradling, 1997; Nakamura et al., 2001; Tritschler et al., 2008; Wang et al., 2017; Wilhelm et al., 2003). Interestingly, following the rapid destruction of ME31B, TRAL, and Cup, the remaining ME31B (at 10-to 20-fold lower abundance) appears to mediate mRNA destruction, rather than translational repression (Wang et al., 2017). The degradation of ME31B, TRAL, and Cup tantalizingly coincides with many of the hallmarks of the MZT, but explorations into this issue have been hindered by a lack of understanding of how their destruction is controlled.

We previously made an intriguing observation that genetically linked the clearance of ME31B, TRAL, and Cup, to the Pan Gu (PNG) kinase (Wang et al., 2017). Composed of three subunits (PNG, Giant Nuclei [GNU], and Plutonium [PLU]), the PNG kinase is central to the oocyte-to-egg transition and mediates key aspects of embryogenesis, including resumption of the cell cycle, zygotic transcription, and maternal mRNA clearance (Elfring et al., 1997; Tadros et al., 2007; Vardy and Orr-Weaver, 2007). Unlike many animals, the oocyte-to-egg transition in *Drosophila* does not require fertilization but is instead triggered by egg activation (Doane, 1960; Heifetz et al., 2001; Horner and Wolfner, 2008a). Here, the PNG kinase is activated by mechanical stress as the oocyte passes through the oviduct, and then phosphorylation and degradation of the GNU subunit quickly inactivates the kinase, restricting its activity to the first half hour after egg activation (Hara et al., 2017). One way that PNG mediates the oocyte-to-embryo transition is by rewiring post-transcriptional gene regulation (Eichhorn et al., 2016; Kronja et al., 2014). Possibly by phosphorylating key RNA binding proteins such as Pumilio, PNG changes the poly(A)-tail length and translation of thousands of transcripts during egg activation (Hara et al., 2018). Importantly, two targets induced by PNG activity are the pioneer transcription factor Zelda, which is responsible for ZGA, and the RNA binding protein Smaug, which is responsible for clearance of many maternal transcripts (Benoit et al., 2009; Eichhorn et al., 2016; Liang et al., 2008; Tadros et al., 2007; Vardy and Orr-Weaver, 2007). The PNG kinase also phosphorylates ME31B, Cup, and TRAL (Hara et al., 2018), but it is unclear what effect phosphorylation has on these proteins. One possibility has been that PNG phosphorylation could lead to the degradation of ME31B, TRAL, and Cup, but this model has been thus far unexplored.

The ubiquitin-proteasome system is a major protein degradation pathway. Here, a series of ubiquitin activating enzymes, conjugating enzymes, and ligases (E1, E2, and E3, respectively) lead to the post-translational addition of a polyubiquitin chain to a target protein, which then serves as a molecular beacon for degradation by the proteasome. E3 ligases are typically thought to recognize target proteins, while E2 conjugating enzymes provide the activated ubiquitin and in turn recognize the E3 ligase (Komander and Rape, 2012). There are hundreds of different E3 ligases and 29 annotated E2 conjugating enzymes in *Drosophila* (Du et al., 2011), but most of the client substrates are unknown, and surprisingly few have been implicated in the MZT.

Given the key roles of ME31B, Cup, and TRAL in oogenesis and embryogenesis, we wanted to understand the mechanisms controlling their degradation. In particular, we sought to answer how PNG activity at egg activation leads to the degradation of these three RNA binding proteins several hours later, and how their degradation is coordinated with other elements of the MZT, including ZGA and maternal mRNA clearance. To answer these questions, we performed a small selective RNAi screen in *Drosophila*, and identified the E2 conjugating enzyme as UBC-E2H/Kondo and the E3 ligase as the CTLH complex. Importantly, the CTLH complex recognized and bound ME31B even in the absence of PNG activity, indicating that phosphorylation does not trigger destruction of this protein. In contrast, *Kondo* mRNA is translationally upregulated by more than 20-fold upon egg activation in a PNG-dependent manner. Thus, egg activation through PNG starts a timer for ME31B, Cup, and TRAL destruction, mediated by the translational upregulation of *Kondo*.

## RESULTS

### PNG kinase activity at egg activation triggers destruction of ME31B

We previously demonstrated through western blotting that ME31B, TRAL, and Cup were degraded 2-3 hours after egg laying (Wang et al., 2017). To understand the mechanisms underlying degradation of these RNA binding proteins, we decided to establish a fluorescence-based assay so that we could follow ME31B degradation in living embryos. To do so, we took advantage of an ME31B-GFP trap line where the fusion protein is expressed from the endogenous locus (Buszczak et al., 2007); we have previously shown that ME31B-GFP recapitulates the dynamics of the wild-type protein (Wang et al., 2017). Consistent with western blotting, GFP signal in *png^50^*/FM7 embryos robustly decreased from 2 to 3 hours after egg laying (Figure 1A). In contrast, the GFP signal remained constant through this time period in *png^50^*/*png^50^* embryos (hereafter referred to as *png^50^*). These results confirm that the differences in ME31B dynamics are observable by microscopy and that the degradation of ME31B requires PNG.

**Figure 1.**
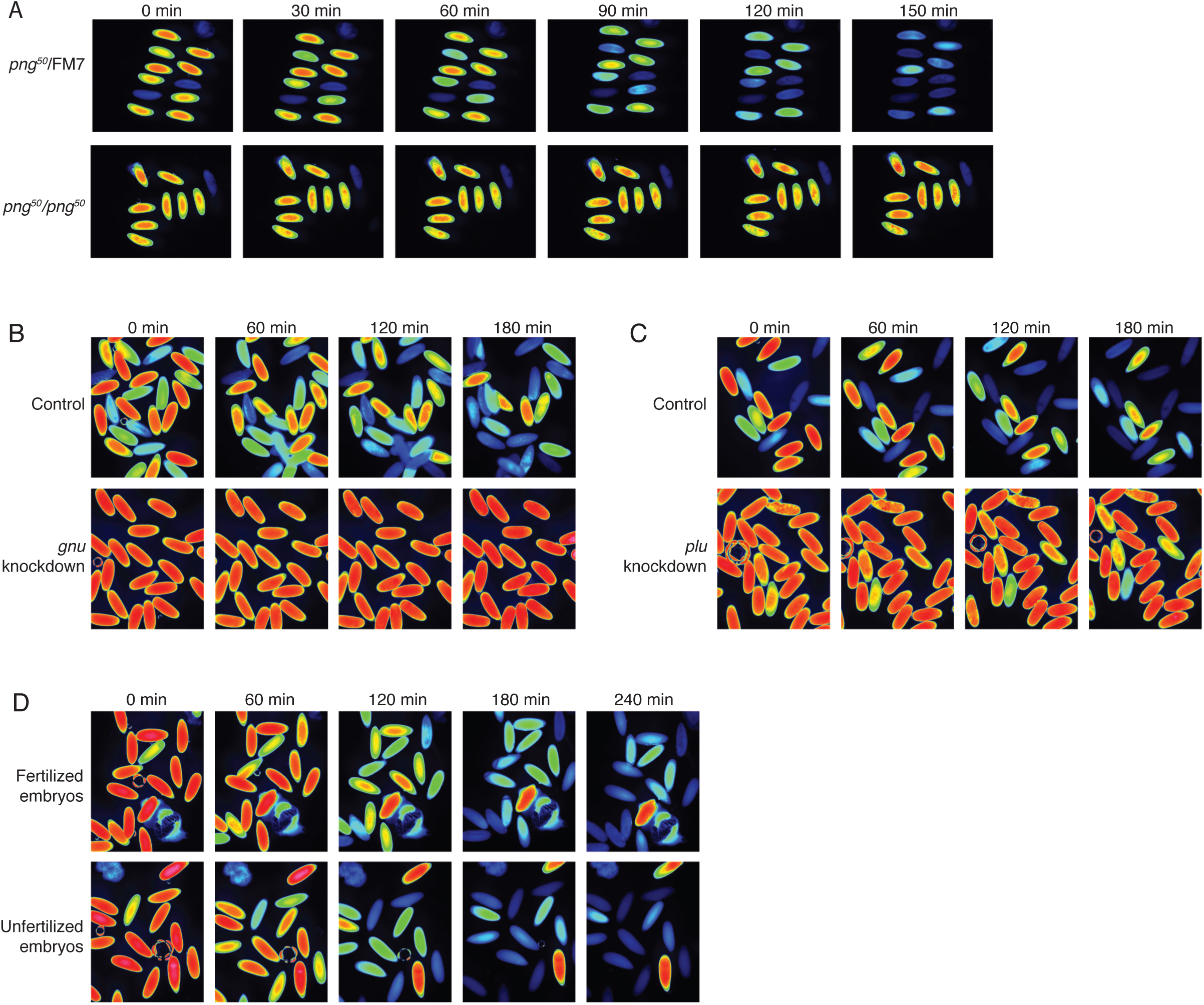
The PNG kinase, but not fertilization, is required for ME31B degradation. (A) PNG is required for the destruction of ME31B. Embryos from female flies of the indicated genotypes (that also contained the ME31B-GFP allele) were visualized at various time points after egg laying. Fluorescent images are false-colored so that more intense fluorescence is indicated by hotter colors. (B) GNU is required for the destruction of ME31B. dsRNA targeting GNU or mCherry (as a control) mRNA was expressed during oogenesis; female flies also contained the ME31B-GFP allele. Laid embryos were visualized by fluorescence microscopy at indicated time points. (C) PLU is required for the destruction of ME31B. As in B, except with dsRNA targeting PLU mRNA. (D) Fertilization is not required for the destruction of ME31B. As in B, except with fertilized and unfertilized embryos.

To test the importance of other genes for ME31B degradation, we combined the ME31B-GFP allele into a GAL4-UAS system, where GAL4 was under the control of the matα-tubulin promoter and so is specifically expressed during oogenesis; this system enabled us to induce the expression of dsRNA during oogenesis and monitor the requirement of various genes for ME31B degradation. Given that *png^50^* embryos did not degrade ME31B, we first investigated whether the other two components of the PNG kinase, GNU and PLU, were required. When either GNU or PLU were knocked down (Figure 1B, C), ME31B was again stabilized, thus confirming that its destruction requires the full PNG kinase.

We next asked whether the degradation of ME31B required fertilization. To do so, we obtained unfertilized embryos, and again followed ME31B levels with fluorescence microscopy (Figure 1D). In contrast to our results in *png^50^* embryos, ME31B was still unstable in the activated, but unfertilized, eggs. This result is consistent with numerous studies demonstrating that the major events pre-MZT in *Drosophila* require egg activation (primarily through the PNG kinase), but not fertilization (Fenger et al., 2000; Horner and Wolfner, 2008b; Tadros et al., 2007; 2003; Vardy and Orr-Weaver, 2007). Taken together, we conclude that degradation of ME31B, and likely Cup and TRAL, is triggered by egg activation through PNG activity.

### ME31B is degraded by the ubiquitin-proteasome system

We hypothesized that ME31B degradation involved ubiquitination. To test this model, we immunoprecipitated ME31B-GFP from 1–2 hour lysates under stringent conditions that disrupted protein-protein interactions, such as that with eIF4E, and probed for ubiquitin by western blotting (Figure 2A, B). Consistent with our hypothesis, this experiment demonstrated that ME31B is ubiquitinated in the early embryo (Figure 2A). We also detected ubiquitin by mass spectrometry of ME31B-GFP pull-downs (see below). Furthermore, this ubiquitination was not detectable in *png^50^* mutant embryos and thus depended upon the PNG kinase activity (Figure 2B).

**Figure 2.**
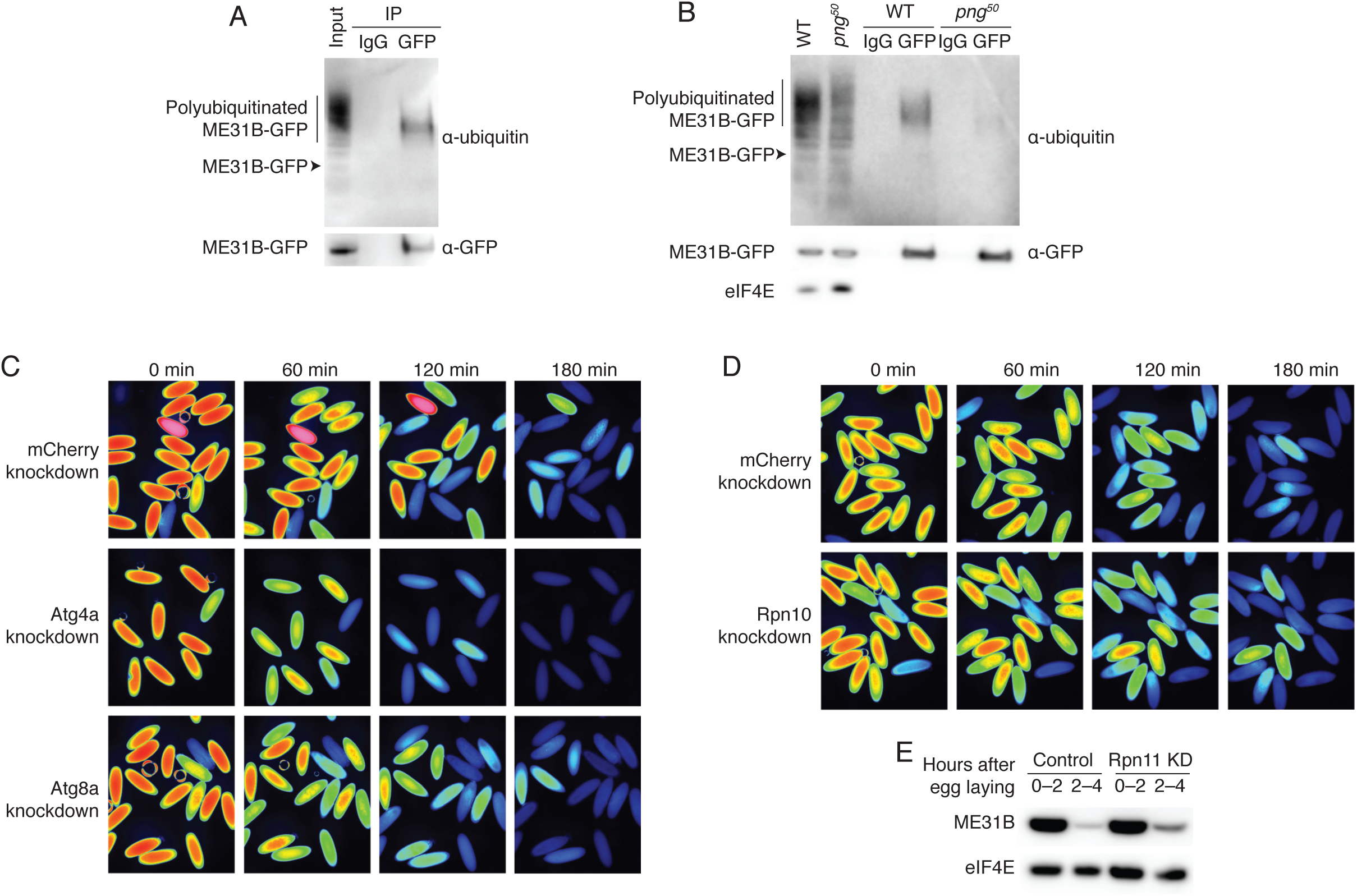
ME31B is degraded by the ubiquitin-proteasome pathway. (A) ME31B-GFP is ubiquitinated. ME31B-GFP was immunoprecipitated from 1–2 hour wild-type embryo lysates under stringent conditions and then analyzed by western blotting by probing with α-GFP or α-ubiquitin. IgG was used as an immunoprecipitation control. (B) Ubiquitination of ME31B-GFP requires the PNG kinase. As in A, except that the immunoprecipitations were performed from wild-type or png50 embryos, and then analyzed by western blotting by probing with α-GFP, α-eIF4E or α-ubiquitin. (C) ME31B-GFP is not degraded by autophagy. Embryos from female flies expressing the indicated dsRNAs and ME31B-GFP were imaged at various time points by fluorescence microscopy. Fluorescent images are false-colored so that more intense fluorescence is indicated by hotter colors. (D) Inhibiting the proteasome partially stabilizes ME31B-GFP. As in C, except for dsRNAs targeting mCherry or Rpn10, a proteasome component. (E) Depleting the proteasome partially stabilizes ME31B-GFP. Staged embryos from the indicated times were harvested with mCherry or Rpn11 knocked down. Western blotting was performed on the lysates, probing for GFP and eIF4E (as a loading control). Related to Table S1.

We next probed whether ME31B is degraded by autophagy or by the proteasome. To do so, we depleted components of either system using the GAL4-UAS system described above. Knockdown of five autophagy components, such as Atg4a or Atg8a, gave viable embryos. However, ME31B was not stabilized in any of these knockdown embryos; indeed, in some cases, it appeared to be degraded more quickly than in the control embryos (Figure 2C, Table S1).

Analysis of embryos depleted of core barrel proteasome proteins proved more challenging because knockdown of most components, such as Prosα5 and Prosα7, resulted in females that did not lay eggs (Table S1). We were able to obtain embryos from Rpn10 and Rpn11 knockdowns, two components of the regulatory particle of the proteasome, likely because these embryos only had partial inhibition of proteasome function or functional redundancy. Importantly, we observed partial stabilization of ME31B in both knockdown embryos (Figure 2D, E). Thus, taken together, these data indicate that the ubiquitin-proteasome system degrades ME31B.

### The E2 conjugating enzyme UBC-E2H is required for the degradation of ME31B, TRAL, and Cup

We thus set out to identify E3 ligases responsible for the degradation of ME31B by carrying out a medium-scale RNAi screen. As before, we monitored ME31B decay by GFP fluorescence, taking images every 30 minutes after egg laying. We focused on those E3 ligases that: (1) had evidence of expression, based on RNA-seq or mass spectrometry data, and (2) had available RNAi lines. We screened 137 RNAi lines, targeting E3 ligases as well as related factors that are involved in sumoylation and neddylation (Table S1). Note that many cullins and proteasomal components fail to lay eggs, presumably because of critical functions during oogenesis.

Our initial expectation was that we would identify one or two complexes required for the degradation of ME31B, but we instead found that factors from many different complexes all appeared partially required, where ME31B degradation was delayed in the knockdown embryos, but eventually proceeded. Hits included multiple components of the SCF E3 ligase (SkpA, Slmb, and Roc1a), both Gus and Cul5 (which work together (Kugler et al., 2010)), and both EloB and EloC. We also found individual components of other E3 complexes that were required, including Poe and the F-box protein Fbw5 (Table S1). Interestingly, knockdown of multiple components of the sumoylation pathway also resulted in slow degradation, including *smt3* (the gene encoding SUMO), both components of the SUMO-activating complex (*Uba2* and *Aos1*), and *degringolade* (*dgrn*), which is the sole ubiquitin E3 ligase in *Drosophila* that recognizes sumoylated proteins (Abed et al., 2011). Given previous reports that ME31B, TRAL, and Cup are sumoylated (Nie et al., 2009), our data suggest that SUMO modification may be important for the degradation of these RNA binding proteins during the MZT.

But, given the incomplete stabilization of ME31B, we hypothesized that there might be another, as-yet unidentified factor important for ME31B degradation, and so we continued the RNAi screen, this time looking at E2 conjugating enzymes. Strikingly, knockdown of UBC-E2H, an E2 ligase conserved from yeast to humans (Kaiser et al., 1995; 1994; Lampert et al., 2018), completely blocked degradation of ME31B, nearly phenocopying the dynamics seen in *png^50^* mutants, as determined by fluorescence microscopy (Figure 3A).

**Figure 3.**
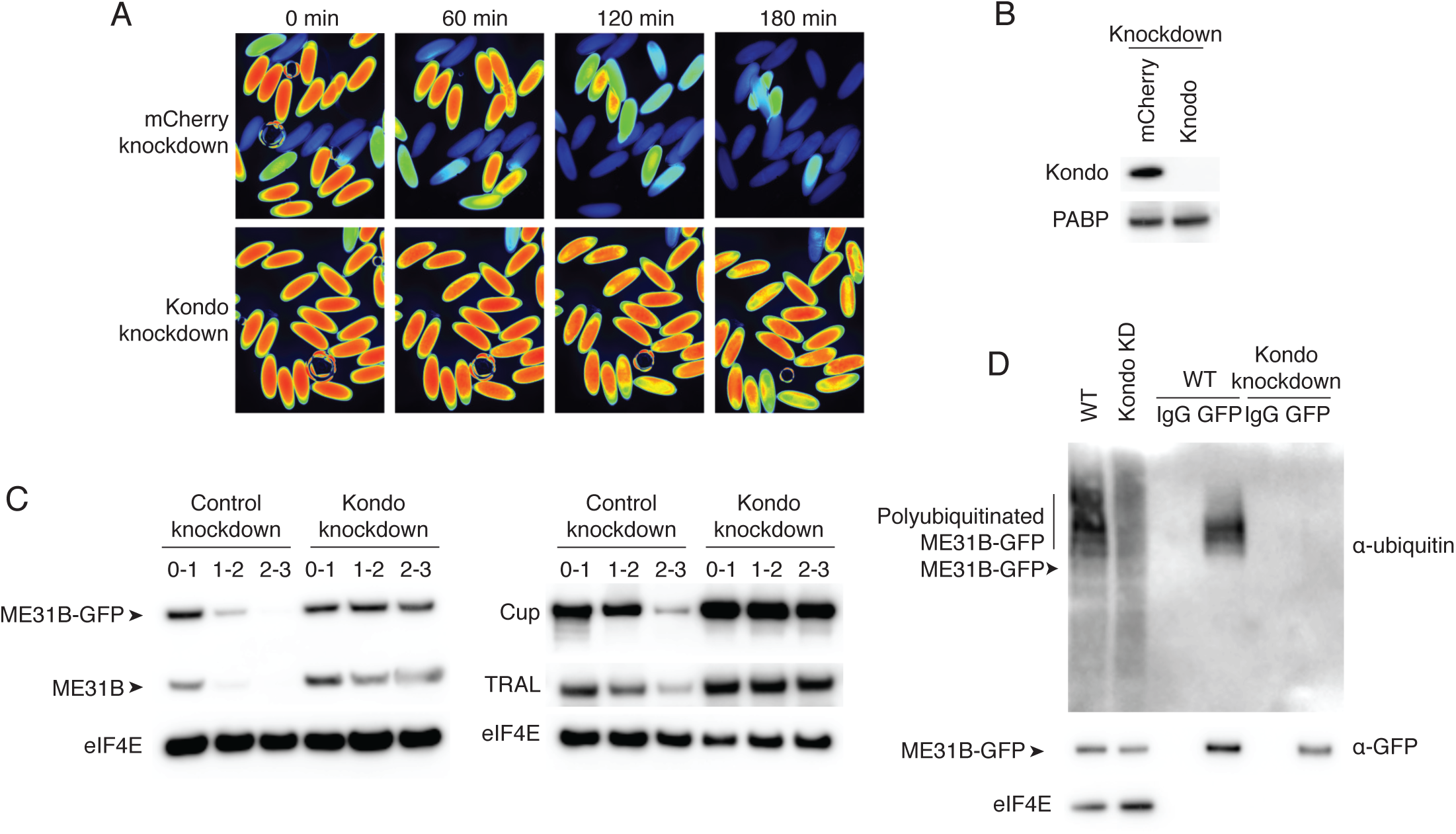
The Kondo E2 conjugating enzyme mediates the degradation of ME31B, TRAL, and Cup. (A) Kondo (UBC-E2H) is required for the degradation of ME31B-GFP. Embryos laid from female flies expressing the indicated dsRNAs and ME31B-GFP were imaged at various time points by fluorescence microscopy. Fluorescent images are false-colored so that more intense fluorescence is indicated by hotter colors. (B) Kondo is depleted in knock-down lines. An antibody was raised against Kondo, and 1–2 hour embryo lysates with the indicated dsRNA were probed for Kondo and PABP (as a loading control). (C) Kondo is required for destruction of ME31B, TRAL, and Cup. Staged embryos with mCherry or Kondo knocked down were harvested at the indicated times. Western blotting was performed on the lysates, probing for ME31B, Cup, TRAL, and eIF4E (as a loading control). Because the maternal flies contained wild-type and ME31B-GFP alleles, both proteins are visible in the α-ME31B blot. (D) Kondo is required for the ubiquitination of ME31B. ME31B-GFP was immunoprecipitated from 1–2 hour mCherry or Kondo knock-down embryo lysates under stringent conditions and then analyzed by western blotting by probing with α-GFP, α-eIF4E, or α-ubiquitin. IgG was used as an immunoprecipitation control. Related to Figure S1 and Table S1.

To test this result, we first raised an antibody against UBC-E2H and confirmed that it was depleted in the knockdown embryos (Figure 3B). We next isolated staged lysates from these embryos and performed western blotting, probing for ME31B. Because the maternal line contains both the wild-type and trap ME31B alleles, this experiment revealed that both wild-type and GFP fusion proteins were completely stabilized when UBC-E2H was depleted (Figure 3C). Moreover, as determined by western blotting, Cup and TRAL were also stabilized in the UBC-E2H knockdown embryos (Figure 3C). Due to its role in removing proteins given as part of the maternal dowry, we renamed UBC-E2H as “Marie Kondo” (shortened to “Kondo”). Finally, through immunoprecipitation experiments, we determined that ubiquitination of ME31B was completely absent in *Kondo* knockdown embryos (Figure 3D). Thus, we conclude that Kondo is required for the ubiquitination and destruction of ME31B, TRAL, and Cup during the MZT.

### Degradation of ME31B, TRAL, and Cup requires the CTLH E3 ligase

Kondo is conserved from yeast to humans and is known to work through the CTLH E3 ligase, a multicomponent complex (Lampert et al., 2018; Santt et al., 2008). Using BLAST, we were easily able to identify putative homologs of the CTLH components in *Drosophila*: RanBPM (homologous to *Hs* RanBP9), Muskelin, CG3295 (homologous to *Hs* RMND5A/GID2), CG7611 (homologous to *Hs* WCR26), CG6617 (homologous to *Hs* TWA1/GID8), and CG31357 (homologous to *Hs* MAEA) (Figure 4A). We were unable to find putative homologs for *Hs* GID5/ARCM8 or *Hs* GID4. Notably, none of these genes were annotated as putative E3 components, and thus none were included in our original screen.

**Figure 4.**
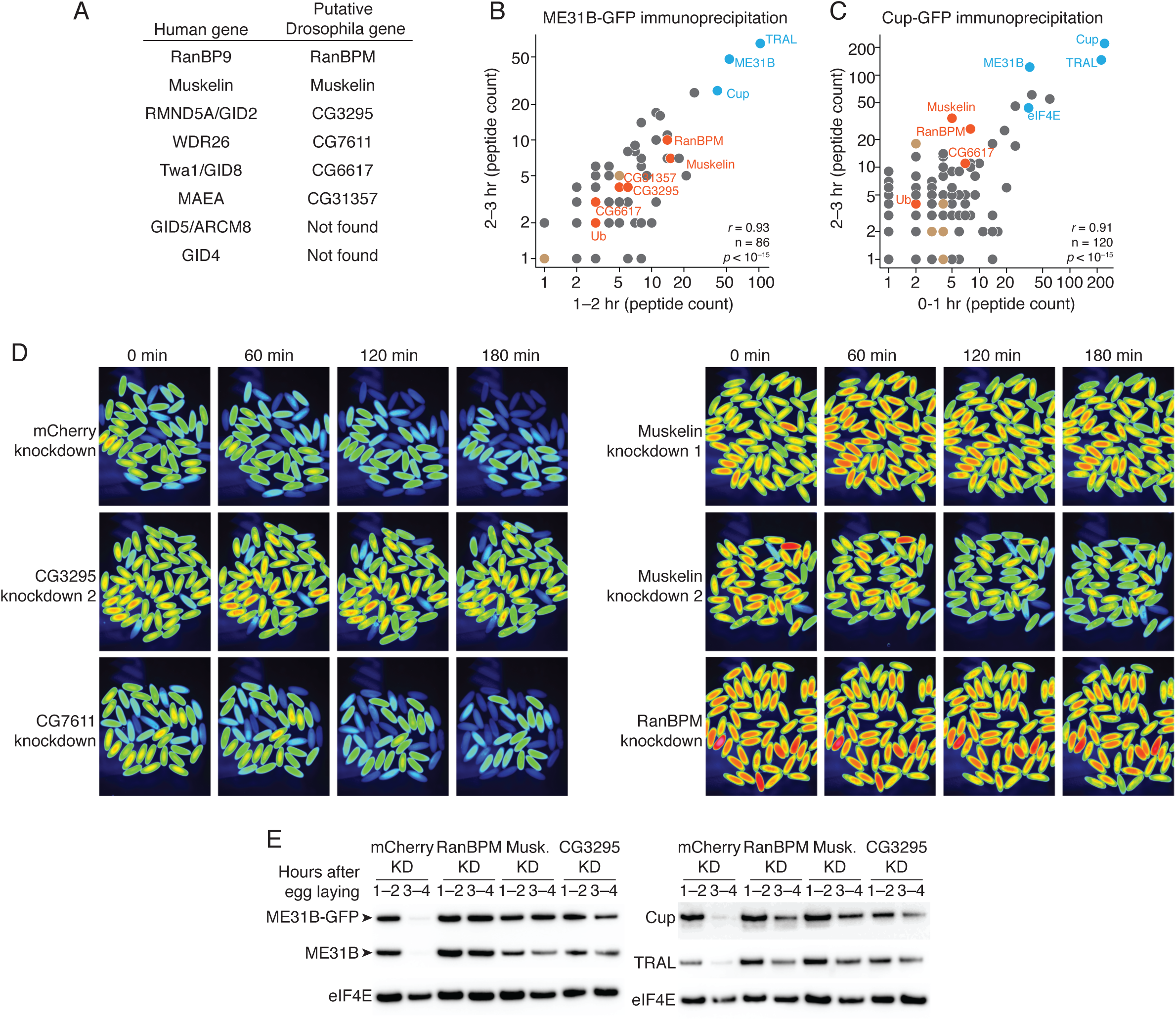
The CTLH E3 ligase mediates the degradation of ME31B, TRAL, and Cup. (A) Putative components of the Drosophila CTLH complex. Drosophila orthologs of the human components of the CTLH complex were determined by BLAST. Some components, such as GID4, appear to not have an identifiable ortholog. (B) ME31B interacts with the CTLH complex. Proteins interacting with ME31B-GFP at 1–2 and 2–3 hour after egg laying were determined by immunoprecipitation with α-GFP and mass spectrometry. Plotted are number of peptides detected (after subtracting for the signal in the control IgG immunoprecipitation). Known interactors with ME31B, such as Cup and TRAL, are shown in blue; CTLH components and ubiquitin are shown in orange; other hits from the screen are shown in brown. (C) Cup interacts with the CTLH complex. As in B, except for Cup-GFP. (D) The CTLH complex is required for destruction of ME31B-GFP. Embryos laid from the indicated maternal RNAi lines were visualized for three hours after egg-laying for ME31B-GFP fluorescence. Fluorescent images are false-colored so that more intense fluorescence is indicated by hotter colors. (E) The CTLH complex is required for destruction of ME31B, TRAL, and Cup. Staged embryos from the indicated times were harvested with mCherry, RanBPM, Muskelin (Musk.), or CG3295 knocked down. Western blotting was performed on the lysates, probing for ME31B, Cup, TRAL, and eIF4E (as a loading control). Because the maternal flies contained wild-type and ME31B-GFP alleles, both proteins are visible in the α-ME31B blot. Related to Figure S2.

To ask if the CTLH complex might be involved in the degradation of ME31B, we immunoprecipitated ME31B-GFP and Cup-GFP from pre-MZT embryos (in conditions that maintain complex formation) and determined the proteins bound by mass spectrometry (Figure 4B, 4C). Consistent with our previous work (Wang et al., 2017), both immunoprecipitations readily pulled-down other members of the Cup–TRAL– ME31B complex. This analysis identified ubiquitin, components of the SCF complex (both SkpA and Slmb), Gus, and both EloB and EloC, suggesting that these factors directly or indirectly interact with ME31B and Cup. Strikingly, however, components of CTLH were even more abundant than these other E3 ligases, and we were able to identify Muskelin, RanBPM, and CG6617 in all four samples. We also detected CG3295 in both ME31B pull-downs and one Cup pull-down, and CG31357 in both ME31B pull-downs. Similar results were seen in previous studies of ME31B complexes in embryonic lysates (Götze et al., 2017). We were also unable to detect CG7611 in any of our samples, despite its ortholog being a component of the human CTLH complex.

We next asked whether destruction of ME31B required the CTLH E3 ligase. As before, we knocked down various components using available RNAi lines (CG3295, CG7611, Muskelin, and RanBPM), and monitored levels of ME31B by fluorescence microscopy (Figure 4D). In contrast to what we had observed with the previous E3 ligases, depletion of CG3295, Muskelin, or RanBPM almost completely stabilized ME31B. Notably, depletion of CG7611, the one component that we failed to detect by mass spectrometry, did not affect the destruction of ME31B. Moreover, when we used western blotting to look at levels of wild-type ME31B, Cup and TRAL, we found that all of these proteins were stabilized when the CTLH complex was depleted (Figure 4E).

One trivial explanation for these results is that depletion of the CTLH complex inadvertently reduced levels of Kondo, which is required for the destruction of ME31B (Figure 3). However, as determined by western blotting, Kondo levels were unaffected in these knockdown embryos (Figure S2A). Because antibodies were only available for RanBPM (Dansereau and Lasko, 2008), we were unable to generally determine how depletion of individual components affected levels of the other components. Nonetheless, in analyzing RanBPM levels, we unexpectedly found that the RanBPM RNAi line only depleted one of its two isoforms, thus suggesting that only the long RanBPM isoform is required for the destruction of ME31B (Figure S2B). Taken together, we conclude that the CTLH E3 ligase is required for the destruction of ME31B, Cup, and TRAL, and, in the early *Drosophila* embryo, is likely composed of RanBPM, Muskelin, CG6617, CG3295, and CG31357. Because of their roles in clearing proteins, we also now refer to CG6617 as Houki (Hou; Japanese for “broom”), CG3295 as Souji (Sou; Japanese for “cleaning”), and as CG31357 as Katazuke (Kaz, Japanese for “tidying up”).

### Association between the CTLH E3 ligase and ME31B does not require PNG kinase activity

Having identified the E3 ligase and E2 conjugating enzyme required for the destruction of ME31B, we next turned to understanding how its destruction was triggered by PNG activity. Because recent work has demonstrated that the PNG kinase phosphorylates ME31B, TRAL, and Cup (Hara et al., 2018), we explored the idea that this phosphorylation might stimulate an association between ME31B and the E3 ligase.

We first asked whether we could detect an interaction between ME31B and the CTLH complex using immunoprecipitation followed by western blotting. Consistent with our mass spectrometry analysis (Figure 4A), we detected RanBPM in ME31B immunoprecipitations (Figure 5A). Interestingly, only the long isoform of RanBPM interacted with ME31B, consistent with this (but not the shorter) isoform being required for the degradation of ME31B, Cup, and TRAL (Figure 4E, 5A). The interaction between RanBPM and ME31B depended upon Muskelin, but not Souji (Figure 5B), although much about the organization of the CTLH complex remains unknown. Moreover, consistent with a role for the E3 ligase binding a target protein, interaction between RanBPM and ME31B was unaffected by Kondo depletion (Figure 5C). Strikingly, however, the interaction between RanBPM and ME31B remained robust in *png^50^* embryos (Figure 5D). Thus, we conclude that PNG activity (and thus phosphorylation of ME31B) is not required for the CTLH complex to recognize and interact with ME31B.

**Figure 5.**
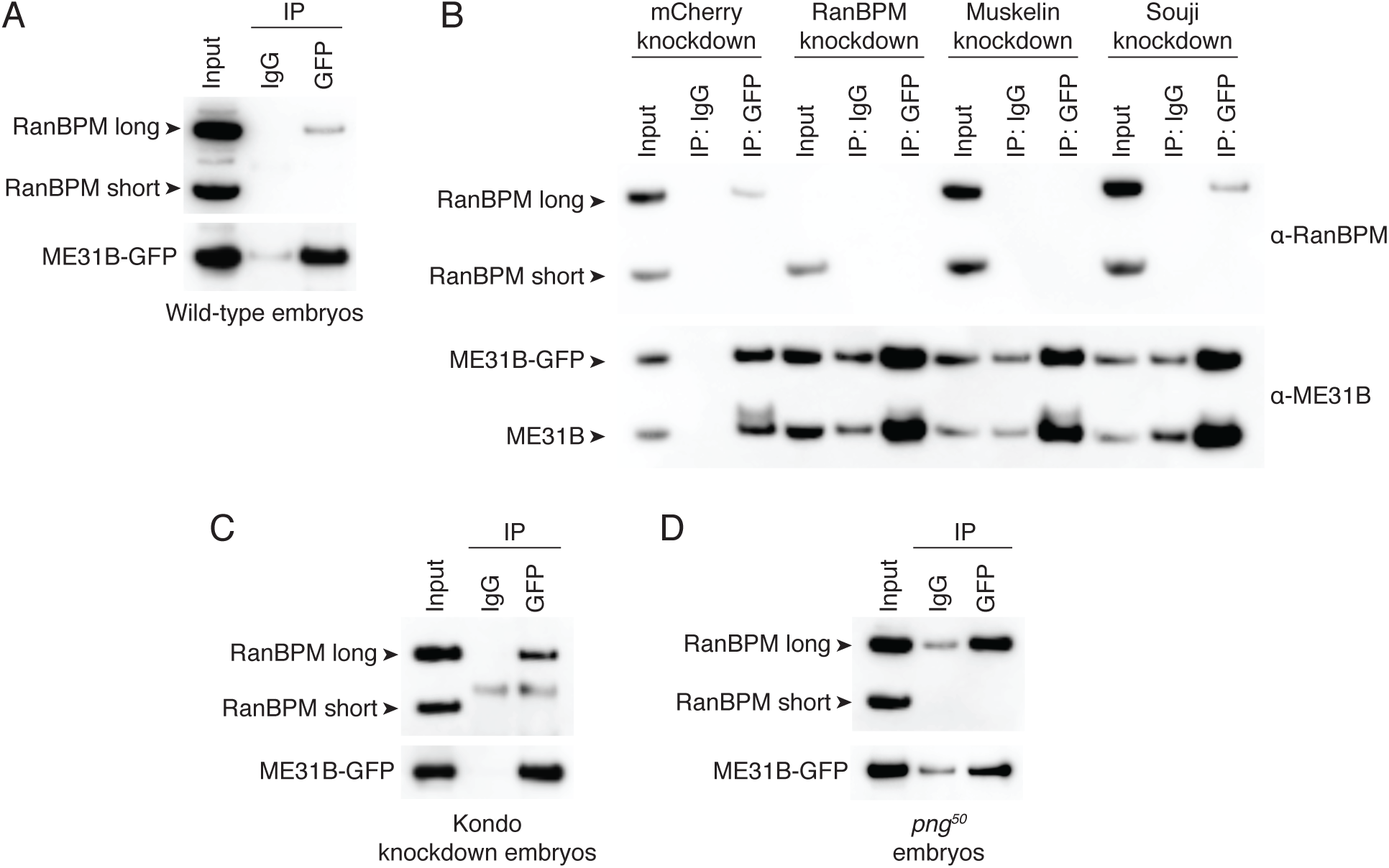
The interaction between ME31B and the CTLH complex does not depend on PNG. (A) ME31B-GFP interacts with RanBPM, a component of the CTLH complex. Proteins interacting with ME31B-GFP at 1–2 hour after egg laying were determined by immunoprecipitation with α-GFP or IgG (as a control) and western blotting. Membranes were probed with α-GFP or α-RanBPM. (B) The interaction between ME31B-GFP and RanBPM depends on Muskelin. As in A, except immunoprecipitations were performed in 1–2 hour embryo lysates with mCherry, RanBPM, Muskelin, or CG3295 knocked-down. (C) The interaction between ME31B-GFP and RanBPM does not depend on Kondo. As in B, except in 1–2 hour embryo lysates with Kondo knocked-down. (D) The interaction between ME31B-GFP and RanBPM does not depend on PNG. As in B, except in 1–2 hour *png^50^* embryo lysates.

### Egg activation mediates the translational upregulation of Kondo via the PNG kinase

Given that PNG phosphorylation of ME31B could not explain how egg activation triggered its destruction, we searched for alternative explanations, focusing on recent observations that PNG also mediates the translational upregulation of thousands of mRNAs at the oocyte-to-embryo transition (Eichhorn et al., 2016). We analyzed published ribosome profiling datasets (Eichhorn et al., 2016) for evidence of translational upregulation of CTLH components and *Kondo* mRNAs upon the oocyte-to-embryo transition. The CTLH component mRNAs were either not affected or downregulated during egg activation (Figure S3). In contrast, the most striking change occurred for translation of *Kondo* mRNA: although translation of *Kondo* mRNA was repressed through oogenesis, its translation increased 25-fold during the oocyte-to-embryo transition (Figure 6A), placing it in the top 10% of genes upregulated at this developmental transition.

**Figure 6.**
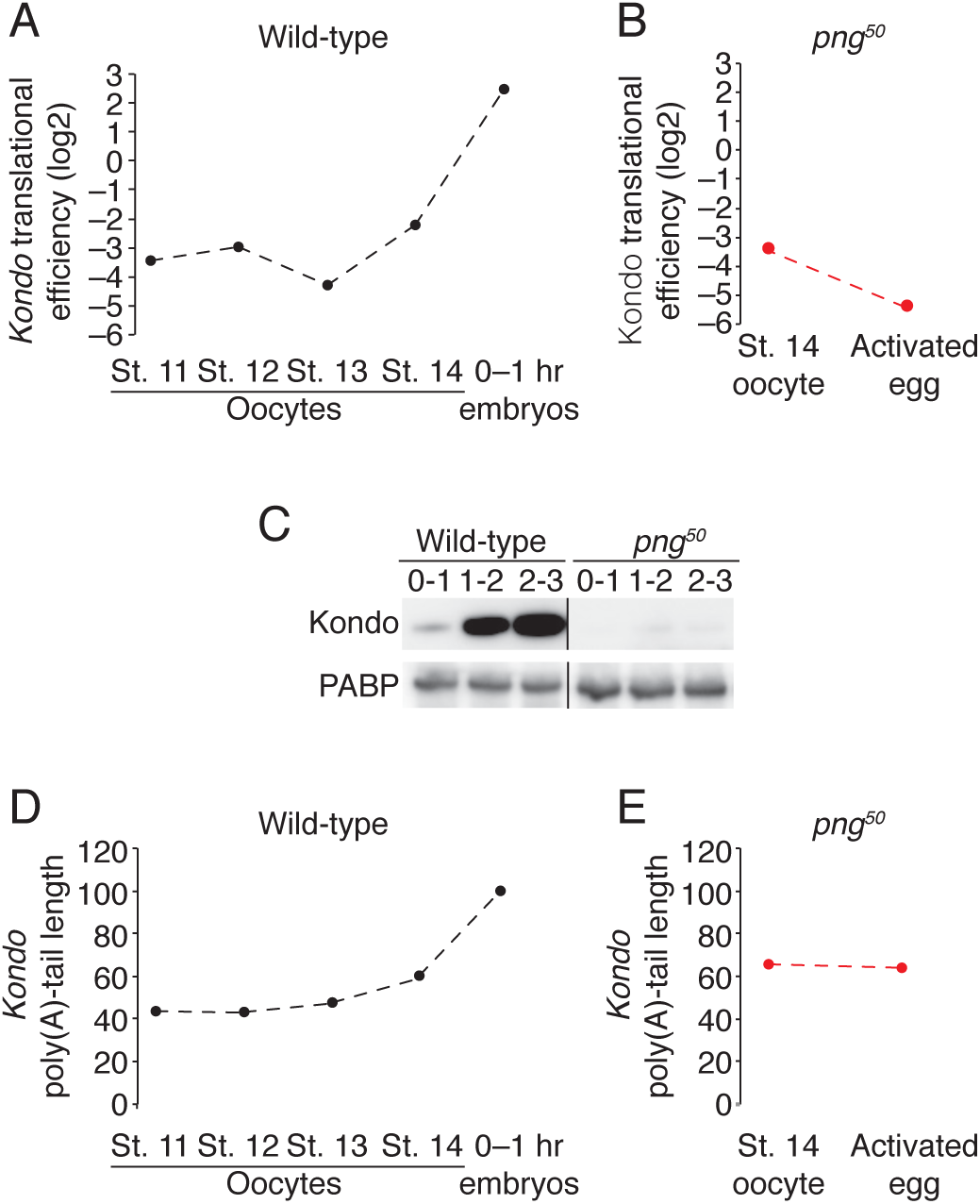
The PNG kinase is required for Kondo production in early embryogenesis. (A) Translation of Kondo increases at egg activation. Translational efficiency of Kondo mRNA was measured in published ribosome profiling datasets (Eichhorn et al., 2016). Shown is the translational efficiency in stage 11, 12, 13, and 14 oocytes, and 0–1 hour embryos. (B) Translational up-regulation of Kondo depends on PNG. As in B, except for *png^50^* stage 14 oocytes and activated eggs. (C) Production of Kondo during embryogenesis depends on PNG. Staged wild-type or png50 embryos were harvested at the indicated times after egg laying. Western blotting was performed on the lysates, probing for Kondo and PABP (as a loading control). (D) Kondo mRNA poly(A)-tail length increases at egg activation. As in A, except plotting poly(A)-tail length. (E) Lengthening of Kondo mRNA poly(A)-tail length depends on PNG. As in B, except plotting poly(A)-tail length. Related to Figure S3.

We next hypothesized that the translational control of Kondo depended upon PNG. We again made use of published ribosome profiling and analyzed the change in translation of *Kondo* mRNA during egg activation in *png^50^* mutants (Figure 6B). While upregulated in wild-type embryos, *Kondo* was not translationally upregulated in *png^50^* activated embryos: its translation differed by more than 200-fold between wild-type and mutant activated eggs. In fact, *Kondo* was the seventh-most affected transcript, showing a similar dependence on PNG as *Smaug*, which encodes a well-known and important downstream target of PNG activity.

Consistent with this analysis, when we probed Kondo protein levels, we were able to detect a dramatic increase in protein levels over the first three hours of embryogenesis in wild-type embryos, but we were unable to detect expression at any time point in *png^50^* mutant embryos (Figure 6C). Pointing to a role of post-transcriptional gene regulation, the poly(A) tail length on *Kondo* mRNA also doubled upon egg activation in a manner dependent on PNG. This result suggests that the translational increase is partially mediated by an increase in poly(A) tail length (Figure 6D, E), which is directly coupled with translation at this developmental stage (Eichhorn et al., 2016). Together, these data have uncovered a genetic network where egg activation turns on the PNG kinase and so leads to the robust translational activation of *Kondo*, which encodes an E2 conjugating enzyme required for the destruction of ME31B, TRAL, and Cup.

## DISCUSSION

ME31B, Cup, and TRAL are RNA binding proteins that are degraded during the MZT, and their destruction coincides with the major wave of zygotic transcription and clearance of maternal mRNAs. Despite occurring several hours after egg laying, degradation of these proteins is triggered by egg activation through the activity of the PNG kinase and is mediated by the ubiquitin-proteasome system (Figure 1, 2). Through a small-scale RNAi screen, we identified that the E2 conjugating enzyme Kondo is required for the ubiquitination and clearance of ME31B, TRAL, and Cup (Figure 3). Kondo is conserved from yeast to humans and, as in those systems (Kaiser et al., 1994; Lampert et al., 2018), appears to work with the CTLH complex, which acts as the E3 ligase. Components of the CTLH complex physically interact with ME31B, and the CTLH complex is also required for the degradation of ME31B, TRAL, and Cup during early embryogenesis (Figure 4). Surprisingly, the association of CTLH with ME31B occurs in the absence of PNG activity, suggesting that, although ME31B (as well as TRAL and Cup) are phosphorylated by the kinase, phosphorylation is not required for their destruction (Figure 5). Instead, translation of *Kondo* is dramatically upregulated at the oocyte-to-embryo transition, in a process that depends on PNG activity (Figure 6). Together, these data suggest a model that egg activation via the PNG kinase leads to translational activation and production of Kondo, which then allows the CTLH complex to ubiquitinate ME31B, TRAL, and Cup and ultimately leads to their destruction (Figure 7). Although the CTLH complex is conserved, it has not yet been studied in *Drosophila*. Our data point to this complex being composed of multiple components (Muskelin, RanBPM, Houki, Souji, and Katazuke), as in other organisms. However, due to a lack of available reagents, we do not know about the stoichiometry of these components, and it remains possible that there are additional, *Drosophila*-specific components. Nonetheless, so far the CTLH complex appears simpler than the human and yeast complexes. We were unable to identify orthologs of GID4 and GID5, which are seen in the human and yeast versions (Lampert et al., 2018; Maitland et al., 2019; Santt et al., 2008). Similarly, although WDR26 is important in the human complex and we identified a *Drosophila* ortholog (CG7611), we found no evidence of its association with ME31B or requirement for ME31B degradation. Thus, we conclude that it is either a nonessential component or not part of the *Drosophila* CTLH complex. Interestingly, the role of the CTLH complex during the MZT appears very different from its role in glucose metabolism (in yeast) and DNA damage (in humans) (Lampert et al., 2018; Santt et al., 2008). In the future, it will be important to determine other proteins targeted by the *Drosophila* CTLH complex and the extent to which ME31B, a ubiquitously expressed protein, is targeted outside of the MZT.

**Figure 7.**
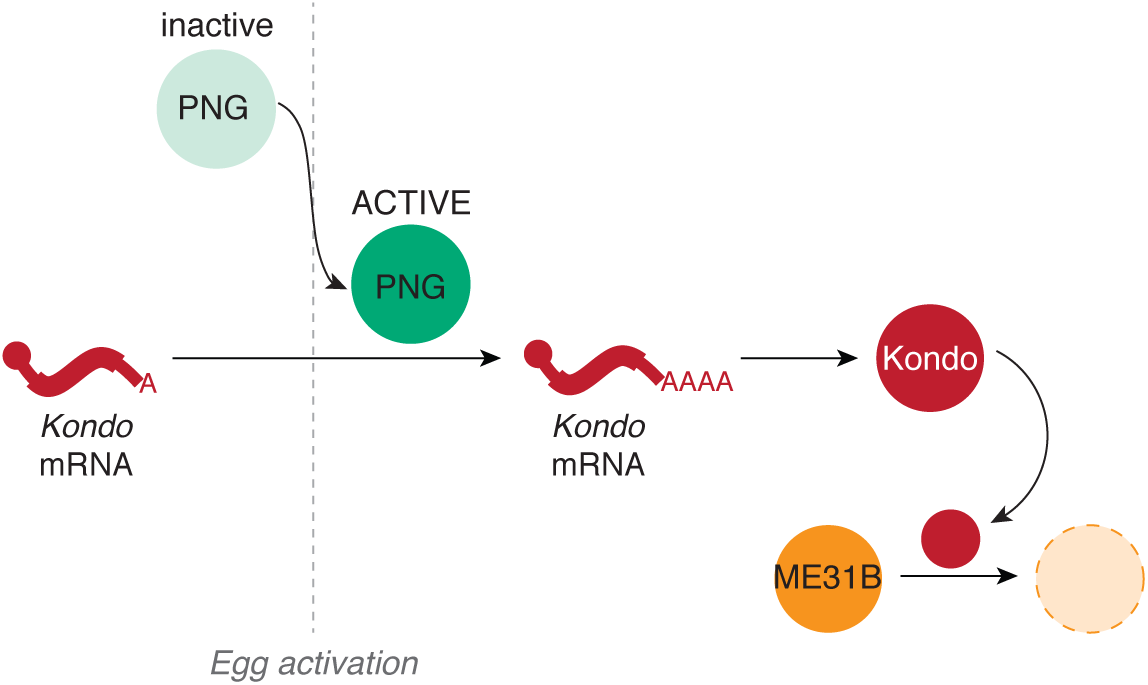
Model for how egg activation results in the clearance of ME31B, TRAL, and Cup. During egg activation, mechanical stress as the egg passes through the oviduct activates the Pan Gu (PNG) kinase. The activated kinase then mediates the translational upregulation of Kondo mRNA, possibly by alleviating repression by 3’UTR-binding proteins. Upon production of Kondo, the CTLH E3 ligase is able to ubiquitinate and destroy ME31B, Cup, and TRAL.

One surprising result is the role of PNG in mediating the destruction of ME31B, TRAL, and Cup. PNG phosphorylates all three proteins (Hara et al., 2018), and so our initial hypothesis was that this modification also stimulates their destruction. However, contrary to our expectations, ME31B interacted with the CTLH complex even in *png^50^* embryos, demonstrating that the phosphorylation is not required for recognition of ME31B by the E3 ligase. An unresolved question, then, is how PNG phosphorylation affects the activities of ME31B, TRAL, and Cup. Intriguing observations from the Orr-Weaver lab suggest that the modification can impact the ability of these proteins to repress gene expression (Hara et al., 2018). It is tempting to speculate that phosphorylation may then contribute to the MZT by modulating the activities of ME31B, TRAL, and Cup, rather than their stability.

The link between PNG and the destruction of ME31B, TRAL, and Cup instead appears to be mediated through the translational upregulation of *Kondo*. Although PNG may contribute through other, as-yet undiscovered, mechanisms as well, this link is sufficient to explain the PNG requirement for ME31B degradation: in the absence of Kondo, ME31B is stable during the MZT, and in the absence of PNG, Kondo is not expressed. An important question for the future will be to understand how PNG mediates the upregulation of *Kondo* translation. One hint may be that the 3’UTR of *Kondo* contains several putative Pumilio binding sites, and translation of *Kondo* is upregulated in ovaries where Pumilio has been knocked down (Flora et al., 2018). Pumilio is also a target of PNG (Hara et al., 2018), and so a possible model is that translational repressors, such as Pumilio, are phosphorylated and inactivated at egg activation, leading to the production of Kondo.

Interestingly, PNG also mediates the translational upregulation of key MZT effectors: Zelda, the pioneer transcription factor, and Smaug, an RNA binding protein that targets nearly two-thirds of the maternal transcriptome for degradation (Chen et al., 2014; Eichhorn et al., 2016; Tadros et al., 2007). Together with our results, a picture is emerging that egg activation stimulates the production of multiple key factors that are important for clearing the maternal RNA and protein dowry and for producing zygotic gene products.

Although the MZT has typically been considered from the perspective of RNA, a role for maternal protein clearance is becoming more important. Over the past few years, the list of proteins degraded during the *Drosophila* MZT has grown and now includes GNU, Matrimony, Cort, Smaug, ME31B, TRAL, and Cup (Benoit et al., 2009; Hara et al., 2017; Pesin and Orr-Weaver, 2007; Wang et al., 2017; Whitfield et al., 2013). Unbiased mass spectrometry experiments also suggest that Wispy and Dhd are also robustly degraded (Sysoev et al., 2016). As this list of proteins in *Drosophila* and other developmental systems increases, a new question is emerging: how many maternally deposited proteins are degraded during the MZT? Understanding the mechanisms controlling protein degradation during the MZT as well as the impact of removing the maternal protein dowry will be key issues to explore in the future.

## MATERIALS AND METHODS

### Drosophila fly stocks

Fly stocks were maintained in a 25°C incubator with 65% humidity.

### Microscopy

Male flies from the TRiP stocks were crossed with ME31B-GFP driver line. Female flies from this cross were collected and crossed with *w1118* males. For egg collection, flies were transferred in the morning to egg-collection chambers on apple juice/agar plates. Flies were allowed to lay eggs for 1 hour. Eggs were collected into cell strainers, washed with 1X PBS, dechorionated with 25% bleach, and washed with 1X PBS. Dechorionated eggs were transferred onto a glass slide and covered with halocarbon oil 700 (Sigma). Images were taken on a ZEISS SteREO Discovery.V8 microscope with X-Cite 120Q fluorescence illumination system (Exelitas Technologies).

### Isolation of embryos

Embryos were collected at various time points post-egg laying, dechorionated with bleach, and washed with 0.1% Triton X-100. Embryos were then homogenized in lysis buffer B (150 mM KCl, 20 mM HEPES-KOH pH 7.4, 1 mM MgCl_2_, 1mM DTT, complete mini EDTA-free protease inhibitors), and were clarified at 15,000 rpm, 4°C for 15 minutes. The supernatant was stored at −80.

### Western blotting

The rabbit anti-Kondo antibody was generated by Pacific Immunology. For western blotting, it was used at 1:10,000. Rat anti-Cup (a gift of C. Smibert) was used at 1:5,000. Mouse anti-ME31B antibody (a gift of K. Nakamura) was used at 1:5,000. Rabbit anti-TRAL (a gift of K. Nakamura) was used at 1:5,000. Rabbit anti-eIF4E (a gift of E. Izaurralde) was used at 1:10,000. Rabbit anti-PABP (a gift of E. Izaurralde) was used at 1:10,000. Rabbit anti-RanBPM (a gift of P. Lasko) was used at 1:10,000. Mouse anti-GFP (Roche) was used at 1:1,000. Mouse anti-ubiquitin (ThermoFisher Scientific) was used at 1:5,000.

### Immunoprecipitations

For GFP-ME31B immunoprecipitations, pre-made lysates were diluted to 1.0 mg/ml, and then incubated with anti-GFP (Roche) or rabbit IgG (Abcam) for 1 hour, rotating at 4°C. EZView protein G affinity beads (Sigma) were washed 3X with lysis buffer, and 25 μl of slurry was added to the lysate-antibody mixture and incubated for 1 hour, rotating at 4°C. Beads were washed three times with lysis buffer. For western blot analysis, the beads were boiled in loading sample buffer and reducing agent, and immunoprecipitates were loaded onto an SDS-PAGE gel. For RanBPM and ME31B, 2.4% input and 17% IP were loaded.

For immunoprecipitations to test for ubiquitination, pre-made lysates were incubated at 4°C with anti-GFP or rabbit IgG for 1 hour, rotating. EZView protein G affinity beads were washed 2X with RIPA buffer, and then 25 μl of the slurry was added to the lysate-antibody mixture. The lysate-antibody mixture with the beads was diluted with RIPA buffer supplemented with 50 μM PR-619 and protease inhibitors, then incubated for another hour at 4°C, rotating. After incubation, the beads were washed three times with supplemented RIPA buffer and transferred to a new tube. Beads were boiled in loading sample buffer and reducing agent, and immunoprecipitates were loaded on an SDS-PAGE gel. To probe for ubiquitin, 3% input and 20% IP were loaded; to probe for GFP, 1% input and 10% IP were loaded; to probe for eIF4E, 3% input and 10% IP were loaded.

### Tandem mass spectrometry

Immunoprecipitates were sent to SPARC BioCentre (SickKids) for LC/MS/MS analysis.

## ACKNOWLEDGEMENTS

We thank Dr. Julie Claycomb, Dr. Suja Jagannathan, Dr. Howard Lipshitz, Dr. Tania Reis, and members of the Rissland lab for helpful and thoughtful discussions. Support was provided from the University of Colorado RNA Bioscience Initiative (OSR) and NIH grant R35GM128680 (OSR).

---

**Figure S1.**
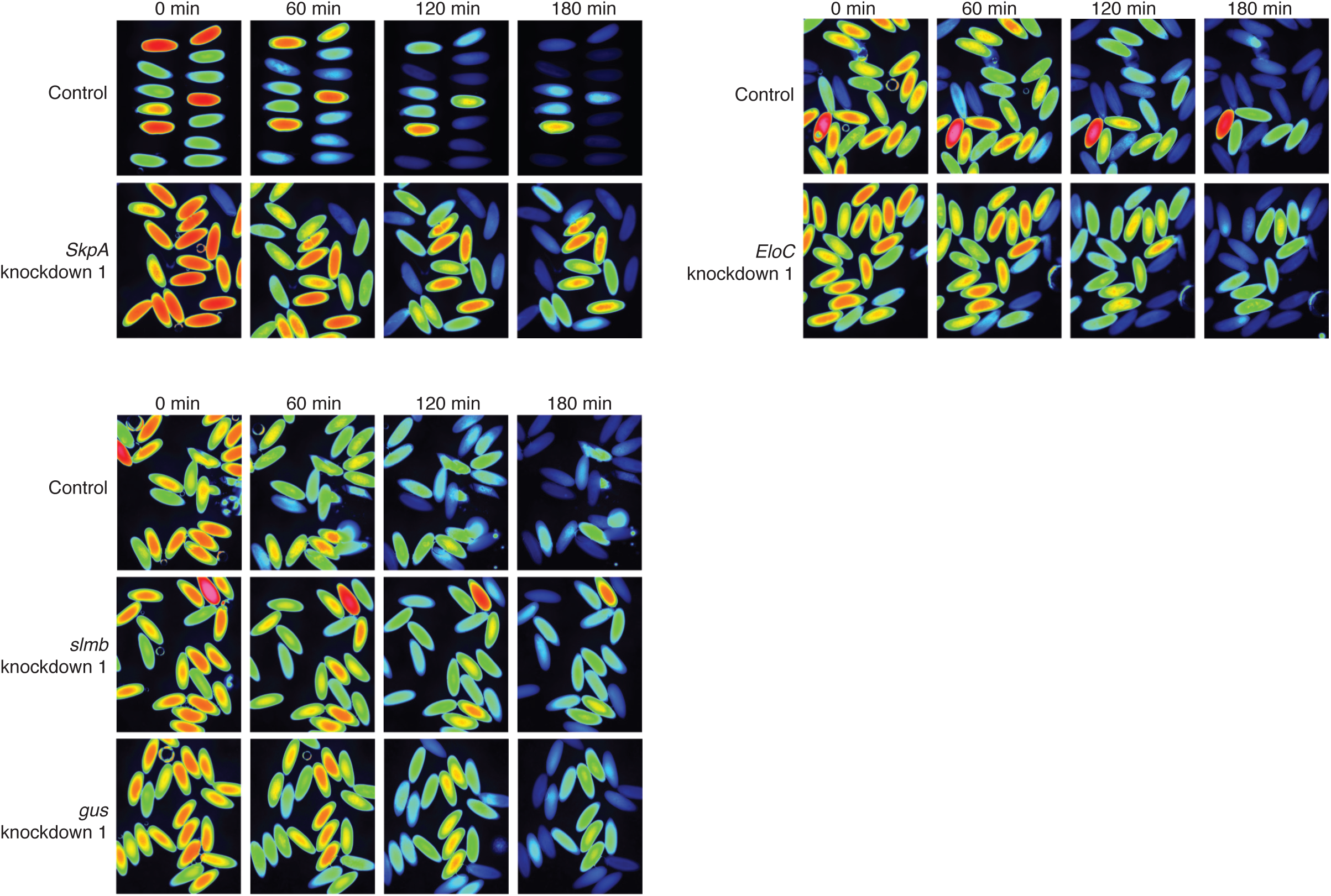
Depletion of several E3 ligases partially stabilizes ME31B-GFP. Embryos laid from female flies expressing the indicated dsRNAs and ME31B-GFP were imaged at various time points by fluorescence microscopy. Fluorescent images are false-colored so that more intense fluorescence is indicated by hotter colors.

**Figure S2.**
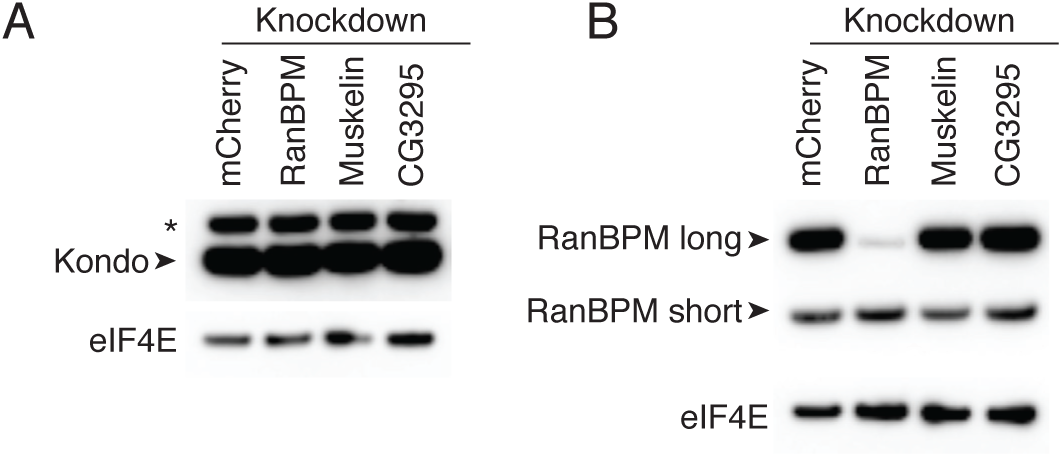
Depletion of CTLH components does not impact Kondo levels. (A) Kondo levels are not affected by CTLH complex depletion. As in A, except probing for Kondo and eIF4E. * indicates a non-specific band. (B) Depletion of Muskelin and CG3295 does not affect RanBPM levels. Western blotting was performed lysates from 1–2 hour embryos with the indicated knock-downs, probing for RanBPM and eIF4E (as a loading control). There are two isoforms of RanBPM, and only the long form is targeted by the RanBPM RNAi.

**Figure S3.**
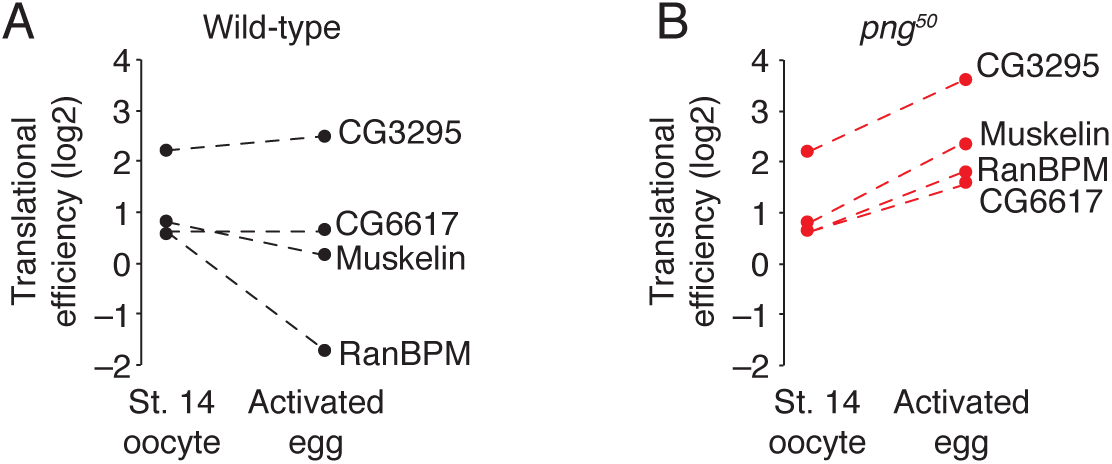
Translation of CTLH components does not increase during the oocyte-to-embryo transition. (A) Translation of CTLH components does not increase at egg activation. Translational efficiency of CTLH components was measured in published ribosome profiling datasets (Eichhorn et al., 2016). Shown is the translational efficiency in stage 14 oocytes and 0–1 hour embryos. (B) Translation of CTLH components is not stimulated by the PNG kinase. As in B, except for *png^50^* stage 14 oocytes and activated eggs.

